# THE MICROANATOMY OF INTRACRANIAL HUMAN THROMBI BY SINGLE NUCLEI TRANSCRIPTOMICS

**DOI:** 10.64898/2026.09.29.755188

**Authors:** Valia P. Leifer, Michael G. Argenziano, Colin P. Sperring, Wenting Zhao, Julia E. Alexander, Aayushi Mahajan, Julia L. Furnari, Corina P. Kotidis, Grace K. Mandigo, Peter D. Canoll, Peter A. Sims, Sean D. Lavine, E. Sander Connolly, Eleonora F. Spinazzi

## Abstract

**Rationale:** Thrombi are fragile, heterogeneous, and difficult to study with available methods, yet understanding their molecular composition and local microenvironment in stroke is critical to elucidating stroke pathophysiology and improving patient outcomes.

**Objective:** We used a novel platform for harvesting and analyzing intracranial thrombi from endovascular thrombectomy for large vessel occlusion to define the cellular and transcriptomic landscape of thrombi.

**Methods and results:** We performed single-nuclei RNA sequencing (snRNA-seq) on three human intracranial thrombi to define their molecular composition and microenvironment. Across 5,143 nuclei, we identified 8 distinct cellular populations, including monocytes and T cells. Monocytes displayed transcriptional changes on gene and regulon levels suggestive of transdifferentiation. We integrated our snRNA-seq data with publicly available single cell RNA-sequencing (scRNA-seq) data from asymptomatic and symptomatic atherosclerotic carotid plaques, and scRNA-seq data from healthy brain vasculature. We identified shared transcriptional programs between carotid and thrombi monocytes along dimensions including inflammation and endothelial activation, supporting the notion that thrombi monocytes undergo transdifferentiation in stroke. We also inferred potential ligand-receptor interactions that may occur at the site and time of occlusion.

**Conclusion:** Our novel application of snRNA-seq to thrombectomy specimens has allowed us to uncover the full diversity of cell types present in thrombi, including low-abundance and novel cell populations, and to provide evidence of cellular plasticity and intercellular communication at the site of occlusion. This detailed definition of the cellular populations involved in stroke will facilitate mapping of novel interventional targets with direct functional relevance and facilitate greater personalization of post-reperfusion therapy.

## Introduction

Acute ischemic stroke (AIS) is the primary cause of disability worldwide [1]. While endovascular thrombectomy (EVT) remains the most effective treatment for recanalization of patients with AIS due to anterior circulation large vessel occlusion [2], roughly 50% of patients remain disabled even after complete recanalization [2]. As the incidence of AIS is projected to rise with population aging, understanding the pathophysiology and molecular underpinnings of reperfusion dynamics and developing therapies to improve outcomes for AIS survivors is of paramount clinical and socioeconomic importance [3].

Therapeutic targeting of mechanisms involved in impaired microvascular reperfusion has been shown to be neuroprotective in animal models [4], [5], [6], [7], [8], [9]. However, oftentimes, therapeutic agents that have conferred cerebral protection in animal models do not yield clinical benefit [7]. Better understanding of the mechanisms of impaired microvascular reperfusion in humans may help explain and overcome the current limitations of revascularization therapies [10].

The composition of intracranial thrombi has gained an increasing amount of interest in recent years and endovascular thrombectomy provides a unique opportunity to harvest thrombectomy specimens for identification of cellular and molecular changes that occur within the area of transient occlusion. To this end, various national and international initiatives have established large-scale thrombus registries [11], [12].

To date, molecular study of human thrombi has been limited to histopathological and bulk sequencing methods. While these methods have revealed that thrombi possess a complex cellular microenvironment, they are limited in their ability to detect low abundance cellular populations and recapitulate the full range of transcriptional programs in thrombi cells [13], [14]. To do so, high-throughput molecular methods, such as single cell RNA-sequencing (scRNA-seq), capable of individually characterizing and profiling multiple cells of the same type in the same state are needed [15]. However, scRNA-seq study of thrombi has been challenging due to the fragile, small, and highly heterogeneous nature of thrombectomy specimens, the complexities of tissue preparation for sequencing, and the unscheduled, emergent nature of their harvesting.

To overcome the challenges posed by currently utilized population and single cell level approaches, we set out to use single nuclei RNA-sequencing (snRNA-seq); a high-throughput technology ideal for working with scarce, complex, and fragile tissues, from which intact and fresh cells are difficult to obtain and dissociate [16], [17], [18].

With snRNA-seq, nuclei can be readily isolated from frozen tissues using a combination of chemical and physical treatments that circumvent issues associated with scRNA-seq isolation methods, including non-uniform or incomplete dissociation of solid tissues into single cells, RNA degradation, and the introduction of technical artifacts (e.g., stress response) that can be triggered during whole cell dissociation methods [16], [17], [19]. Most importantly, snRNA-seq allows for the in-depth characterization of low-abundance cell populations and the unbiased identification of pathways or targets previously obscured using bulk population sequencing methods [16], [17], [18], thereby providing higher resolution into cellular differences and a better understanding of the function of an individual cell in the context of its microenvironment [20], [21].

Here we present the first study to produce a single nuclei transcriptome-based cellular landscape of human intracranial thrombi, comprised of 8 unique populations of immune cells in distinct effector states, and highlight the importance of cellular plasticity and intercellular communication at the site of occlusion. Through integration of our snRNA-seq data with publicly available scRNA-seq data from carotid plaques and healthy brain vasculature, we also contextualize thrombi in carotid atherosclerotic disease and infer potential ligand-receptor interactions that may occur between thrombus cells and the brain vasculature.

Our novel application of snRNA-seq to thrombectomy specimens provides the highest resolution exploration of cellular differences and a best understanding of individual contextualized cell function in the thrombus microenvironment to date. This detailed definition of the cell communities at play will facilitate mapping of novel interventional targets with direct functional relevance and allow for greater personalization of post-reperfusion therapy.

## Methods

### Tissue Removal, Banking, And snRNA-Seq Approach to Thrombectomy Specimen

This work was approved by the Columbia University Irving Medical Center Institutional Review Board before commencing the study. All human tissue was banked under an IRB-approved protocol (IRB-AAAA4666), which permits the distribution of de-identified tissue samples and associated clinical data to researchers for analysis. All thrombectomy specimens were procured from surgeries at Columbia University Irving Medical Center. Standard-of-care interventional techniques were employed to harvest thrombectomy specimens at the time of revascularization as previously described [10], [11], [12], [22], [23]. We routinely use stent retrievers and aspiration catheter systems to retrieve the thrombus. Our established tissue banking protocol has been optimized since 2020 and is seamlessly incorporated into the standard workflow of the mechanical thrombectomy procedure. The tissue is given to a tissue bank technician in the endovascular suite and processed for banking and snRNA-seq. We use a large-scale microfluidic platform for snRNA-seq, which profiles thousands of individual nuclei from multiple specimens in parallel with rapid, high-efficiency cellular/nuclear capture (>50%) at low library construction costs ($0.03 per nucleus) [24]. We have developed experimental and computational pipelines for routine implementation of these technologies to study a variety of neurosurgical specimens. Our data are publicly available (DOI: 10.17632/wt2typ7pzb.1).

We performed pooled 3’-end snRNA-Seq in 96-well plates, which offers improved sensitivity, higher efficiency, and more uniform coverage in full-length PBMC transcriptome profiling [25]. Libraries were prepared per published protocol, Smart-seq2 method, and sequencing (2 x 75 bp) was performed using an Illumina® NextSeq® [26].

### Unsupervised Clustering, Differential Expression, and Visualization

After snRNA-seq preprocessing, raw count matrices for each thrombus (n=3) were loaded into R (v3.12) for individual quality control and data processing. Across all thrombi, nuclei RNA reads expressed in <3 nuclei were excluded. Seurat objects were then created using Seurat (v5.0.1), before additional filtering to exclude nuclei with <100-250 unique features or >2500 unique features [27]. Thrombi were then integrated using Seurat’s integration protocol, which includes correction for batch effects. Integration anchors were identified using 20 dimensions and 1500 features, and then scaled and principal component analysis was performed using 30 principal components and 20 dimensions. UMAP analysis was performed using 20 dimensions, followed by nearest neighbor graph construction with PCA, 20 dimensions, and k=50. Initial clustering was performed at a resolution of 2.0, after which low quality and/or contaminated clusters were identified and excluded based on low unique features per nuclei, low RNA counts, and ribosomal content. Remaining nuclei (n=5,143) were reclustered at a resolution of 1.0, yielding 8 phenotypically distinct clusters.

Cell identifications were made using top differentially expressed genes and canonical marker genes from the literature, and supported using Azimuth 0.5.0, Seurat’s reference-based mapping pipeline, gene set enrichment analyses (GSEA), gene ontology (GO) analysis, and regulon-level analysis with SCENIC, as described below [28]. Gene set enrichment and ontology analyses for individual cellular populations were conducted using EnrichR 3.2, and Seurat’s integrated EnrichR function, with Wilcoxon Rank Sum testing, a log2 fold-change threshold of 0.5, and a p-value threshold of ≥0.0525 [28]. For thrombus cell-type specific analyses, we used databases available through EnrichR.2, including 2021 Descartes Cell Types and Tissues, Panglao 2021, Reactome 2022, Go Biological Processes 2023, and Wiki Human 2024 databases to identify biologically relevant pathways [28]. We used CellChat 2.1.1 to infer potential intra-thrombi ligand-receptor interactions with a p-value threshold of ≤0.05 [29].

### Single Cell Regulatory Network Analysis

Single cell regulatory network analysis was conducted using SCENIC (v1.3.1) using default settings to determine master regulon and transcription factor regulatory networks [30]. Human cis Target databases were downloaded from https://resources.aertslab.org/cistarget/. We used the GENIE3 algorithm to identify co-expression of transcription factors with their target genes, followed by RcisTarget to identify regulon enrichment, and then AUCell to score the regulon activity in each nucleus [30]. Regulon activity by cell cluster was visualized using a heatmap and dot plot.

### Carotid Plaque Data Integration and Analysis

After cell-type specific identifications and analyses, thrombi snRNA-seq data were integrated with publicly available data from asymptomatic and symptomatic carotid plaques (GEO: 155513) using the Seurat integration protocol, with parameters including: integration anchors (20 dimensions, 1500 anchors), principal components (n=30, 20 dimensions), UMAP analysis (PCA reduction, 20 dimensions), nearest neighbor graph construction (PCA reduction, 20 dimensions, k=50), and clustering (resolution 0.6) [31]. Diffusion coefficient analysis (DCA) was subsequently performed to infer changes in transcriptional programming across monocytes from asymptomatic and symptomatic carotid plaques and all three thrombi using the Bioconductor package Destiny 3.1 [32]. For DCA, the Seurat object was first converted to a SingleCellExperiment object using the SingleCellExperiment package version 1.24.0 [33]. The SingleCellExperiment object was then processed for diffusion coefficient analysis using a local Gaussian kernel and 20 eigenvectors. Plotly version 4.10.4 was used to construct two- and three-dimensional visualizations. For comparative gene set enrichment analyses, we used EnrichR 3.2, as well as Seurat’s integrated EnrichR function, with Wilcoxon Rank sum testing, a log2-fold change threshold of 0.5, and a p-value threshold of p<0.05, and the Go Biological Process 2023 database [28].

### Brain Vasculature Data Integration and Analysis

Thrombi snRNA-seq data were integrated with scRNA-seq data from healthy brain vasculature (GEO: 173731) using the Seurat integration protocol, with parameters including: integration anchors (20 dimensions, 1500 anchors), principal components (n=30, 20 dimensions), UMAP analysis (PCA reduction, 20 dimensions), nearest neighbor graph construction (PCA reduction, 20 dimensions, k=50), and clustering (resolution 0.6) [34]. Integration anchors were identified using 20 dimensions and 1500 features. Ligand receptor analysis to infer potential signaling interactions between thrombi and the vasculature was performed using CellChat 2.1.1, with a p-value threshold of ≥0.05 [29].

## Results and Discussion

### Single-nuclei RNA-sequencing of Intracranial Thrombi Reveals the Presence of 8 Functionally Distinct Populations of Immune Cells

Across 5,143 nuclei from 3 thrombi, we identified two broad categories of immune cells, monocytes (CD14+) (78%), and T cells (CD6+) (22%) (**Fig 1B**). Canonical gene cluster-based analysis (**Fig 1C**), differential gene expression analysis via Wilcoxon Rank Sum testing (**Fig 1D**), and gene set enrichment analyses (**Fig 1E, Supp Fig 1**), identified 6 distinct monocytes subpopulations: inflammatory monocytes (*IL1B+ IL15+*) (“IMo”), endothelial monocytes (*VEGFA+ PECAM1+ HK2+ ANGPTL4+*) (“EMo”), regulatory monocytes (*IL6R+ CD22+*) (“RMo”), macrophage-derived monocytes (*CD163+ CD68+ CD84+ ALCAM+*) (“MoM”), and two populations of monocytes with neuroepithelial- and microglia-like phenotypes, one of which displayed increased expression of inflammatory genes (*IL1B+ IL15+ RNASET2+ TYROBP+*) (“NMoI”), while the other expressed neurotransmitter- and inflammation-related regulatory factors, (*GABARAP+ NRGN+ PRNP+*) (“NMoR”). Both T cell subpopulations predominantly consisted of CD4+ cells, including a population of Th2 cells (*GATA3+ STAT6+ IL5+ IL6+*) (“Th2”) and a population of Th17 cells (*RORA+ IL6R+ IL21R+*) (“Th17”). Interspersed among these CD4+ T cells was a small number of CD8+ T cells (∼16), too few to form a statistically significant cluster. We used EnrichR to conduct gene ontology, gene set enrichment analyses (GSEAs), and Seurat’s Azimuth reference-based mapping pipeline to support our cell type identifications and illuminate the unique phenotypic properties of each population (**Supp Fig 1,3)**.

**Figure 1.**
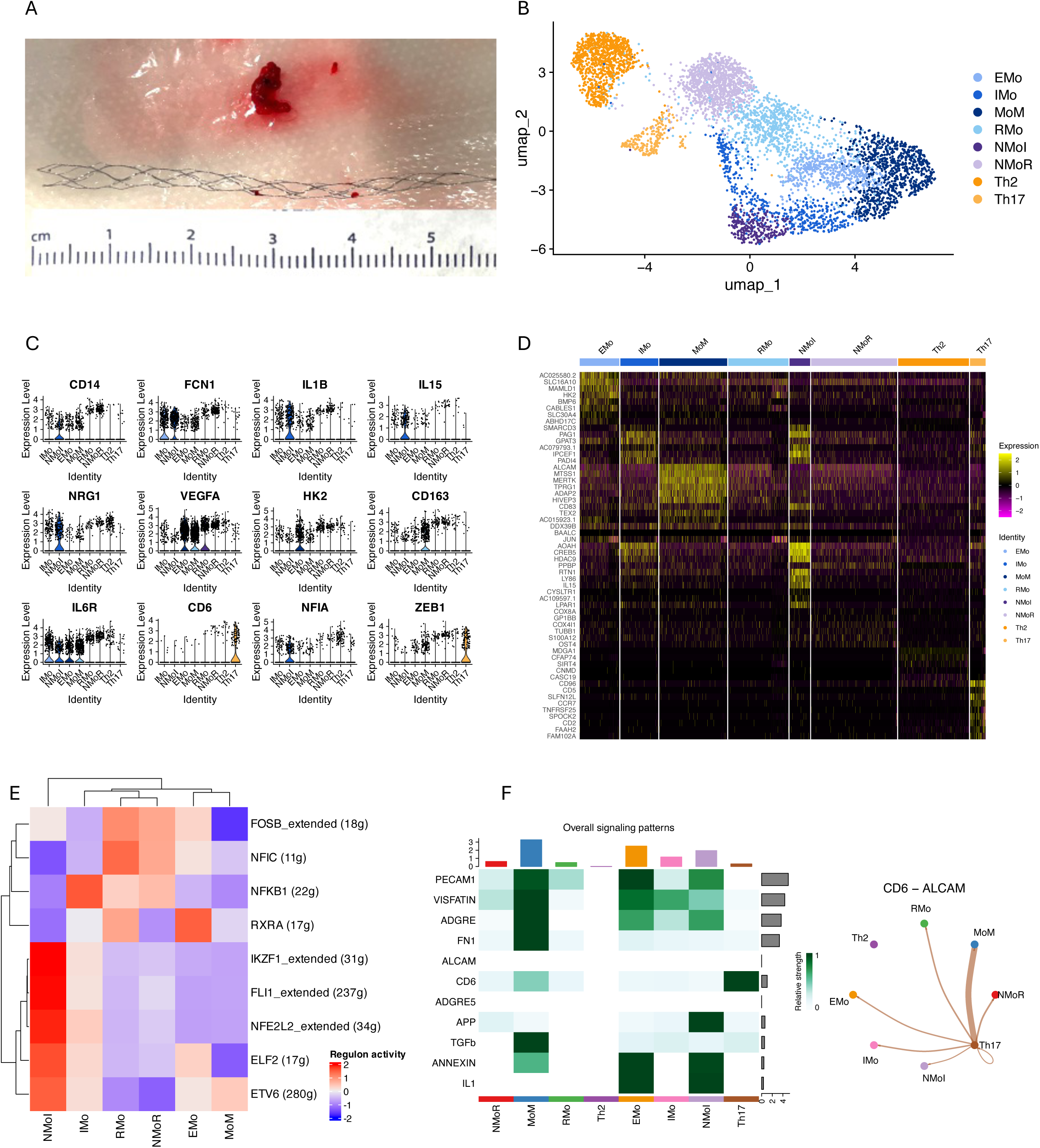
Identification of cellular composition of three intracranial thrombi harvested from endovascular thrombectomy. (A) Intracranial thrombus. Thrombi from endovascular thrombectomy for large vessel occlusion were harvested for single nuclei RNA-sequencing. (B) Uniform manifold approximation and projection analysis plot of 8 phenotypically distinct cellular populations from nuclei of three intracranial human thrombi (n=5,143). (C) Violin plots depicting expression levels of select marker genes used to identify thrombi monocytes/macrophages and T cell populations. (D) Heatmap of top differentially expressed genes by cluster, identified using Wilcoxon rank sum testing, which were used to characterize thrombi cellular populations. Each row represents a gene and each column bar represents a specific cellular population. (E) SCENIC regulon analysis reveals differential regulon activity among thrombi monocyte populations. Each row represents a specific transcription factor and each column represents an individual cluster. The dendrogram at the top represents relatedness among thrombi cell populations based on regulon activity. Activity is relative, with red representing higher levels of activity for a given transcription factor, and blue representing relatively lower levels of activity. (F) Panel 1: Ligand-receptor analysis overview depicting all intra-thrombi communication. Each row represents activity for a specific pathway, and each column represents a specific cell cluster. Panel 2: Ligand-receptor activity of CD6-ALCAM, demonstrating that thrombi monocytes may activate thrombi Th17 cells. EMo, endothelial monocytes; IMo, inflammatory monocytes; MoM, monocyte derived macrophages; RMo, regulatory monocytes; NMoI, neuroepithelial-like inflammatory monocytes; NMoR, neuroepithelial-like regulatory monocytes; Th2, Th2 T cells; Th17, Th17 T cells.

### Two Neuroepithelial-Like Monocyte Clusters Reflect Neurovascular Changes at the Site and at the Time Of LVO

Two neuroepithelial monocyte clusters (NMoI and NMoR) displayed distinct and unexpected gene expression patterns, highlighting the profound influence of the local environment on thrombus monocytes. The neural-specific transcriptional changes observed could only arise from dynamic crosstalk between the thrombus and its surrounding environment. We identified these clusters as monocytes based on their high expression levels of canonical monocyte marker genes, CD14 and CCR2. Given the high degree of phenotypic brain-related change displayed by the NMoI and NMoR clusters, we addressed the possibility that these cells were microglia, rather than embolized monocytes, by evaluating expression levels of microglia marker genes, including IBA1, TMEM19, and P2RY12, by conducting gene set enrichment analyses (GSEAs), and by using Seurat’s Azimuth reference-based pipeline. Across all methods, these clusters emerged as primarily monocytes – both clusters expressed microglia-specific genes at very low to absent levels, and both clusters ranked more highly as monocytes than microglia through GSEA and Azimuth analyses (**Fig 1B-C; Supp Fig 1C-1D, 3**).

### An inflammatory monocyte cluster involved in neuroplasticity and neuroprotection highlights monocyte plasticity and transdifferentiation in stroke

NMoI thrombi monocytes were identified as inflammatory based on high levels of expression of inflammatory cytokine genes such as *IL1B* and *IL15*, and high levels of pathway activity related to immune system activation, including both innate and adaptive pathways. Specific pathway activity included complement activation, neutrophil degranulation, and IL4 and IL13 mediated signaling, which are signatures of the Th2 cells response, consistent with the presence of Th2 cells within the thrombus that we observed (**Supp Fig 1D, 2A**). The finding that monocyte snRNA-seq data can reflect neutrophil-related activity (e.g., degranulation) is notable, given that neutrophils are known to be present in thrombi, but cannot be directly detected using snRNA-seq given that their nuclei are segmented. Finally, GSEA also revealed brain-specific pathway activity suggestive of transdifferentiation in this cluster, including pathways related to complement system in neuronal development and plasticity, HDAC6-related central nervous system activity, neurogenesis, and neuroinflammation.

In addition to inflammatory activity, this cluster also displayed evidence of transdifferentiation related to mitigation of inflammation, including upregulation of genes that are typical of microglia, and the response to inflammation, injury, and hypoxia in the brain. Specific genes included *BTG2*, *ENTPD1*, and *NRG1*. The NMoI cluster’s upregulation of these genes highlights this cluster’s complex transcriptional profile and serves as a reminder that inflammatory monocyte phenotypes exist along a spectrum. *BTG2* is a transcriptional coregulator that is involved in microglia differentiation, and, in preclinical models has been shown to be upregulated in the brain response to hypoxia-ischemia and chronic cerebral hypoperfusion [35], [36], [37]. *ENTPD1* is associated with microglia-dependent suppression of neuronal activity, and in preclinical models has also been shown to promote resistance to occlusive thrombosis after carotid injury ^43,44^. Lastly, *NRG1* belongs to the neuregulin family of proteins, which are structurally similar to epidermal growth factor (EGF) proteins [40]. *NRG1* has been shown to be protective of both brain parenchyma and vasculature in multiple contexts, including neuroinflammation, hypoxia, and excitotoxicity, via suppression of pro-inflammatory cytokines and regulation of neural circuit assembly and synaptic plasticity [40]. In models of ischemic stroke, *NRG1* has even been shown to improve neuronal survival and neurological function [41], [42], [43].

### A neuroepithelial-like cluster involved in neuroprotection and neuroplasticity highlights the potential of snRNA-seq study of thrombi to identify therapeutic targets

NMoR monocytes were identified as regulatory based on expression of genes associated with the transition towards a tissue-resident macrophage phenotype, including *FTH1*, *CD163*, *PF4*, and *TGFB1*, their UMAP association with regulatory monocytes (“RMo”), their overlapping patterns with the RMo cluster in terms of top differentially expressed genes, and GSEA (**Fig 1B-D, Supp Fig 1E, 2B**). Like the NMoI cluster, the NMoR cluster also expressed genes associated with the response to injury, hypoxia, and inflammation in the brain. Notable genes expressed by the NMoR cluster included the GABA type A receptor associated protein (*GABARAP*), neuregulin (*NRGN*), cathepsin S (*CTSS*). In monocytes, *GABARAP* is associated with autophagy and T cell priming; in the brain, it is associated with regulation of GABA-receptor transport to and stabilization at the cell surface/synapse [44], [45]. *NRGN* is a calmodulin-binding protein that is expressed predominantly in the brain, where it is involved in neural plasticity via modulation of calmodulin availability in dendritic spines, and has also been proposed as a novel biomarker of brain injury [46], [47], [48]. Finally, *CTSS* is a lysosomal cysteine protease which is normally secreted by microglia in the brain and has been shown to promote inflammation in stroke [49]. Just as our NMoI cluster did not express only proinflammatory genes, our finding that the NMoR cluster expresses some proinflammatory genes, such as *CTSS*, highlights that thrombi monocytes possess complex, dynamic transcriptional profiles, and that their inflammatory profiles exist along a spectrum.

Together, both clusters’ complex transcriptional profiles raise questions about temporal aspects of transdifferentiation – in each cluster, do monocytes initiate expression of inflammatory and neuroprotective related genes at the same time, or does one phenotypic shift precede the other? The evolution of the expression of pro-vs. anti-inflammatory factors may also be unique to each cluster. Future study involving more thrombi and detailed information about patients’ times to last known normal may help elucidate any potential temporal component to monocyte transdifferentiation in stroke, which may aid not only in predicting patient outcomes, but also in identifying molecular targets with realistic therapeutic windows.

Overall, the NMoI and NMoR clusters highlight the effect that the local environment has on thrombi monocytes in stroke, as these transcriptional changes could only have been induced by the brain. Our findings also suggest that monocytes’ capacity for plasticity, which is well studied in carotid atherosclerosis, is also present in stroke, and can be detected as early as six hours after a patient’s last known normal [31].

### Monocytes enriched in complement and inflammatory factors suggest thrombi may contribute to the onset of inflammation in stroke

We observed the highest degree of complement related activity in the NMoI and IMo clusters, though most thrombi cell populations displayed complement receptor expression (**Supp Fig 5**). Complement is well studied in stroke and of substantial therapeutic interest [50]. Prior work has shown C3a and C5a binding in the lectin and alternative pathways as key effectors of injury; consistent with this, we identified high levels of expression of complement receptor genes *C3R* and *C5R* in our thrombi cell populations. We also identified high expression of *FCN1*, an activator of the complement lectin pathway. The complement activity we observed is also consistent with the presence of the Th17 cells that we observed, as complement can drive Th17 differentiation, and suggests that complement in stroke may promote recruitment and/or activation of this specific T cell type [51], [52]. Altogether, our findings suggest that the thrombus itself may possess the intrinsic ability to initiate complement activity at the site of occlusion.

### The local environment promotes endothelial activation of monocytes in stroke

We also observed monocytes with an endothelial-like phenotype (“EMo”), which were rich in the pro-angiogenic secreted factor *VEGF* and hypoxia-associated genes involving pathways such as glycolysis and gluconeogenesis. This population displayed increased expression of *PECAM1* (*CD31*) and *ITGAX*, which are both endothelial adhesion molecules with pro-angiogenic effects [53]. Upon binding CD99 on endothelial cells, PECAM1 promotes endothelial retraction and monocyte extravasation. While all thrombi monocyte populations displayed increased expression of integrins and selectin-ligands, the EMo population was particularly enriched in markers of endothelial adhesion and pro-angiogenic factors (**Supp Fig 2C**). This finding suggests that the EMo cluster exists in a distinct endothelial-activation state and may carry increased potential to contribute to loss of endothelial integrity and breakdown of the blood brain barrier relative to other thrombi cell populations.

### Transdifferentiation of thrombi immune cells in stroke occurs at both the gene and regulatory network levels

Regulatory gene network analysis of transcriptional expression data using single-cell regulatory network inference and clustering (SCENIC) supported our cluster identifications and aligned with our differential gene expression and gene set pathway enrichment analyses. Our SCENIC analysis supported the notion that the local environment has a dramatic effect on thrombi monocyte and T cell transcriptional dynamics in stroke. Given that gene regulatory networks are identified based on differential expression across clusters, we evaluated regulon dynamics both across all thrombi immune cell populations to highlight regulon-activity among T cells, and across monocytes only to highlight more meaningful regulon-related differences among monocytes. In both analyses, each cluster displayed activation of unique transcriptional programs, with the most dramatic relative changes in upregulated transcription factors occurring in the Th17 and NMoI clusters (**Fig 1E**, **Supp Fig 2**). Based on regulon activity, the NMoI and IMo clusters appeared related, as did the NMoR and RMo clusters (**Fig 1E**).

Regulon analysis comparing monocyte clusters revealed cluster-specific transcription factor activity that related directly to cluster identifications (**Fig 1E)**. The NMoI cluster displayed pathway activity of *IKZF1*, *FLI1*, *NFE2L2*, *ELF2*, and *ETV6*, all of which have been implicated in microglia maturation and/or polarization towards a pro-inflammatory phenotype [54], [55], [56]. *FLI1* and *NFE2L2* have been implicated in microglia differentiation and inflammation [57], [58], [59]. *FLI1* is believed to lead to microglial differentiation and is associated with pericyte loss and neuroinflammation. *NFE2L2* is a CNS- and microglia-specific transcription factor that has been shown to promote antioxidant responses and to be neuroprotective in neurodegenerative diseases; its absence has been implicated in neuroinflammatory disorders such as multiple sclerosis.

Other monocyte clusters displayed upregulation of specific transcription factors also in line with their cell type identifications (**Fig 1E)**. The IMo cluster displayed upregulation of *NFKBI*, a master regulator of inflammation, which is associated with the release of cytokines including TNFa, IL1B, and IL6, as well as inflammasome activation. The RMo and NMoR clusters displayed upregulation of *NFIC*, a positive regulator of monocyte differentiation and promoter of cellular survival, as well as *FosB*, which is induced by hypoxia and induces hypoxia-specific transcriptional programming [60]. The EMo cluster displayed unique activity of *RXRA*, a transcription factor that is directly involved in endothelial activation. *RXRA* has been shown to be neuroprotective in ischemic stroke and also to contribute to polarization of monocytes towards an anti-inflammatory phenotype via inhibition of NF-kB transactivation [61].

Thrombi T cells also displayed unique regulon-level transcriptional programming (**Supp Fig 4**). Th17 cells displayed increased activity of transcription factors related to Th17 differentiation, including *BACH2*, *ZEB1*, and *NFKBI* [62]. *ZEB1*, in particular, is required for pathogenic Th17 differentiation, and also promotes JAK/STAT signaling and interferon-gamma and IL-17 related inflammation [63]. Thrombi Th2 cells displayed increased activity of *NFIA*, also known as *GATA3*, which is a critical transcription factor in Th2 cell differentiation.

### Analysis of intra-thrombi cellular communication reveals that thrombi monocytes may contribute to T cell activation

CellChat analysis revealed potential ligand-receptor signaling interactions among thrombi cell populations (**Fig 1F**). Notably, we observed potential *ALCAM-CD6* activity between macrophage-derived monocytes (“MoM”) and Th17 cells (**Fig 1F, panel 1**). ALCAM-CD6 leads to T cell activation, again suggesting that some degree of post-stroke inflammation may originate, at least in part, from the thrombus itself, and supporting the notion that the thrombus itself is intrinsically inflammatory. Among thrombi cell populations we also identified several interactions related to cellular adhesion and/or migration among monocytes including *PECAM1-PECAM1*, *NAMPT-(ITGA5+ITGB1)*, *(ITGAV+ITGB1)-ADGRE2*, and *ANXA1-FPR1* (**Fig 1F, panel 2**).

### Integration of thrombi single nuclei and carotid single cell data support the notion of monocyte transdifferentiation and interconversion following embolization

To relate thrombi to carotid plaques and contextualize them in the broader picture of atherosclerosis, we integrated our snRNA-seq data with publicly available scRNA-seq data from asymptomatic and symptomatic carotid plaques [31]. In these carotid plaque data, we identified three populations of monocytes (inflammatory carotid monocytes, “cIMo”; non-inflammatory carotid monocytes, “cRMo”; non-inflammatory pro-angiogenic monocytes (“cEMo”); and CD8+ T cells (“cCD8+”)) (**Fig 2A**). Integration of carotid plaque and thrombi data revealed shared transcriptional programming between carotid monocytes and thrombus monocyte/macrophage populations. Of note, T cell populations between the two tissues were not conserved; carotid plaque T cells were all CD8+, while thrombi T cells were mostly CD4+ except for a small number (n=16) of CD8+ cells.

**Figure 2.**
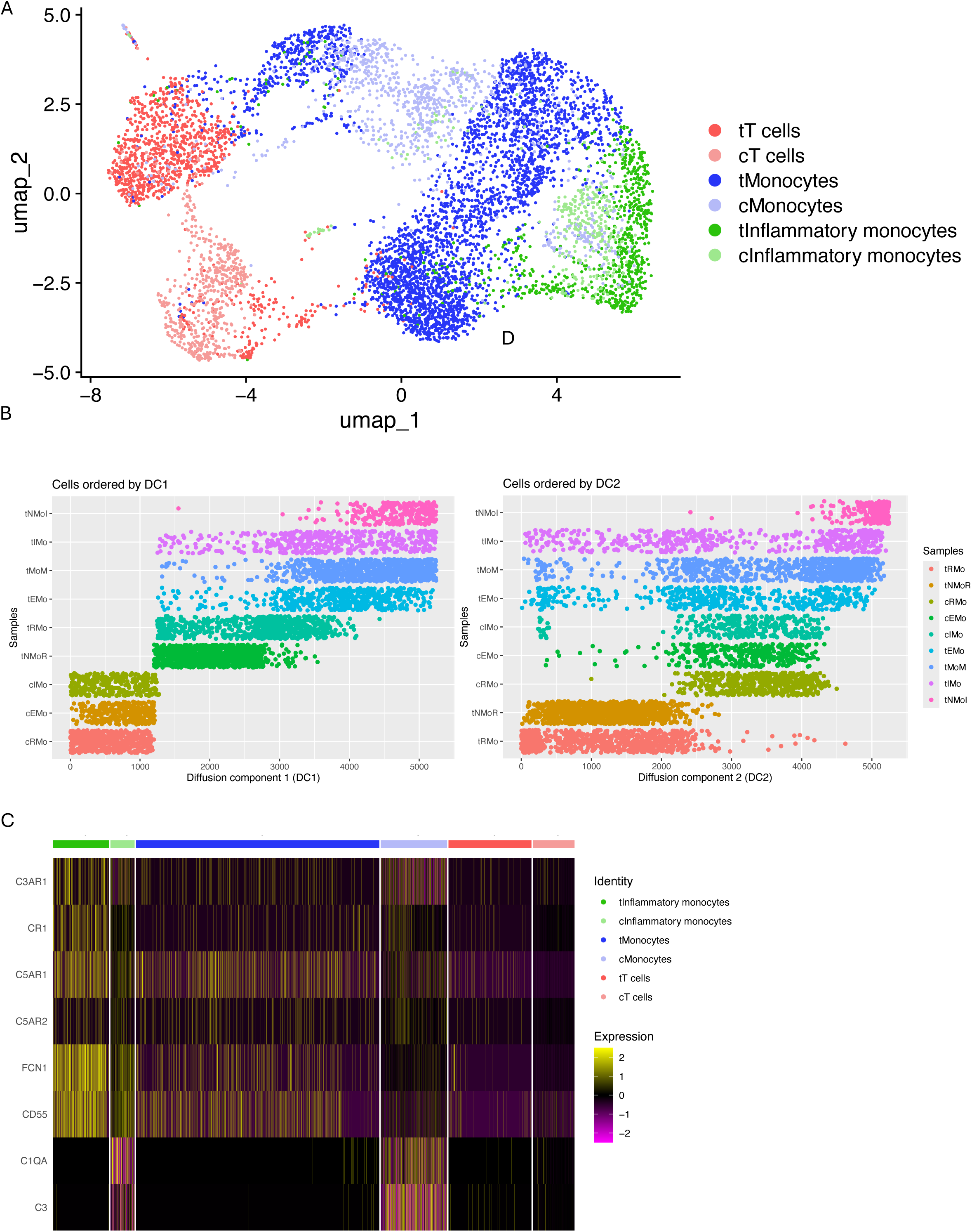
Integration of thrombi snRNA-seq data with publicly available single cell RNA-sequencing data from asymptomatic and symptomatic carotid plaques. (A) Uniform manifold approximation and projection analysis of integrated snRNA-seq and scRNA-seq data from thrombi and carotid plaques. Cells are colored based on broad cell type, T cells, inflammatory monocytes, and non-inflammatory monocytes. “t” indicates thrombi cell populations, “c” indicates carotid cell populations. Thrombi cells are the darker shade for a given broad cell type. (B) Panel 1: Diffusion component (DC) 1 represents inflammation. Here, carotid and thrombi monocytes are displayed along a continuum from the least inflammatory population, carotid regulatory monocytes (“RMo”), to the most inflammatory population, thrombus neuroepithelial-like inflammatory monocytes (“NMoI”). DC1 reveals that even the least inflammatory thrombi monocytes, the neuroepithelial-like regulatory monocytes (“NMoR”), are more inflammatory than the most inflammatory carotid monocyte population (“cIMo”). Panel 2: Diffusion component 2 represents both inflammation and endothelial activation. Here, regulatory thrombus and carotid monocytes fall at one end of DC2, while thrombus monocytes with phenotypes related to inflammation and endothelial activation fall at the other end of DC2. (C) Heatmap depicting relative levels of complement gene expression in thrombi monocytes compared to carotid monocytes. Across each broad category of cell type (e.g., inflammatory thrombi monocytes vs inflammatory carotid monocytes) thrombi cells display higher expression levels of complement related genes. cRMo, carotid regulatory monocytes; cEMo, carotid endothelial monocytes; cIMo, carotid inflammatory monocytes; tNMoR, thrombi neuroepithelial-like regulatory monocytes; tRMo, thrombi regulatory monocytes; tEMo, thrombi endothelial monocytes; tMoM, thrombi monocyte-derived macrophages; tIMo, thrombi inflammatory monocytes; tNMoI, thrombi neuroepithelial-like inflammatory monocytes.

Diffusion coefficient analysis demonstrated that carotid and thrombi monocytes displayed shared transcriptional programs progressing along two primary phenotypic dimensions: inflammation and endothelial activation (**Fig 2B**). Along DC1, which represents the greatest dimension of transcriptional change across all cell populations, both inflammatory and non-inflammatory carotid monocytes sit at one end of the spectrum, while inflammatory thrombi monocytes (NMoI, IMo) sit at the opposite end (**Fig 2B, panel 1**). This finding suggests that DC1 represents both the difference between carotid vs. thrombi cells and inflammation, as the distribution of clusters also progresses from less inflammatory to more inflammatory. In turn, relatively less inflammatory monocyte populations, such as EMo and MoM monocytes, emerge as more inflammatory than inflammatory carotid monocytes.

DC2, which captured the second greatest dimension of transcriptomic change among carotid and thrombi monocyte populations, appeared to represent a combination of endothelial activation and inflammation – here, inflammatory neuroepithelial-like monocytes, inflammatory monocytes, macrophages, and endothelial monocytes fall along one end of DC2, while regulatory monocytes and M2-like macrophages from carotid plaques fall along the other end (**Fig 2B, panel 2**). This interpretation of DC2 was supported using GSEA and gene expression data, as discussed below (**Supp table 1**). Taken together, the DC2 results suggest that endothelial activation is a key dimension of phenotypic change between carotid and thrombi monocytes, which again supports the notion of monocyte plasticity, transdifferentiation, and interconversion following embolization; if this were not the case, and monocytes did not undergo transcriptional reprogramming following embolization, comparison between thrombi and carotid monocyte populations would not reveal phenotypic spectra across diffusion components.

### Comparative analysis of carotid and thrombi monocytes reveals increased complement and inflammatory pathway activity in thrombi

We also evaluated differential complement activity among thrombi and carotid monocytes. Compared to carotid monocytes, thrombi monocytes displayed upregulation of the pro-inflammatory complement component *FCN1*, and complement receptors *CR1*, and *C5R* and *C3R* (**Fig 2C**). While levels of complement factors (e.g., *C1QA, C3*) were low, they were still more highly expressed among thrombi monocytes than carotid monocytes (**Fig 2C**). Of note, thrombi monocytes also displayed expression of *CD55*, an anti-inflammatory negative complement regulator, suggesting a degree of a balance between pro- and anti-complement activity in thrombi, and again highlighting the complex transcriptional profiles these cells possess.

Evaluation of the top 100 differentially expressed pathways (p<0.05) using the Go Biological Process 2023 database between all thrombi and carotid plaque monocytes supported the notion that monocytes progress along inflammatory and endothelial activation-related transcriptional continuums after embolization. Specific pathways upregulated among thrombi monocytes included regulation of vascular associated smooth muscle, regulation of cell death, T cell proliferation, angiogenesis, IL-1-mediated signaling, the response to reactive oxygen species, VEGF signaling, the cellular response to hypoxia, regulation of neuronal morphogenesis and neurogenesis, cytokine-signaling, and leukocyte cell-cell adhesion (**Supp table 1**).

The differences in T cell population identities that we observed between carotid plaques and thrombi deserves further evaluation. In stroke, based largely on animal models, CD8+ T cells are considered the first T cell subtype to arrive at the site of occlusion [64]. While Th2 and Th17 cells have been shown to arrive as early as 24 hours after stroke, their levels are believed to peak 3-7 days after stroke [65], [66]. In stroke, Th2 cells are anti-inflammatory, while Th17 cells are pro-inflammatory [65], [66]. Th17 cells are believed to contribute to the post-ischemic inflammation via secretion of IL-17, which has also been shown to be increased in the peripheral circulation of ischemic stroke patients [67].

While the absence of a statistically significant cluster of CD8+ T cells in our thrombi may be the result of a processing artifact, it is also possible that this finding reflects a true biological process, perhaps either the timing of arrival of distinct T cell subtypes at the site of occlusion, or differences in the relative proportions of T cell subtypes present in stroke in humans compared to animals. Larger scale snRNA-seq based study combined with histopathological staining of sectioned thrombi may help determine whether our observed scarcity of CD8+ T cells is the result of a methodological artifact or reflects a true biological process.

### Brain vasculature integration

To characterize thrombi immune cells and their potential activity in the brain, we integrated snRNA-seq thrombi data with publicly available scRNA-seq data from healthy brain vasculature. Integration revealed that thrombi monocyte subpopulations NMoI and IMo clustered near brain capillary and arterial endothelial cells, that thrombi EMo cells clustered between brain capillary endothelial cells and fibroblasts, and that Th17 cells clustered near pericytes (**Fig 3A,B**). Of note, Th17 cells are able to crawl along pericytes via LFA1, which our Th17 cells expressed, and are believed to contribute to vascular dysfunction and disruption of the blood brain barrier via secretion of IL17, which alters cell-adhesion molecules on endothelial cells [68]. The close phenotypic association between thrombi monocyte populations and endothelial cells in the brain vasculature also supports the notion of transdifferentiation and endothelial activation of these inflammatory cell populations.

**Figure 3.**
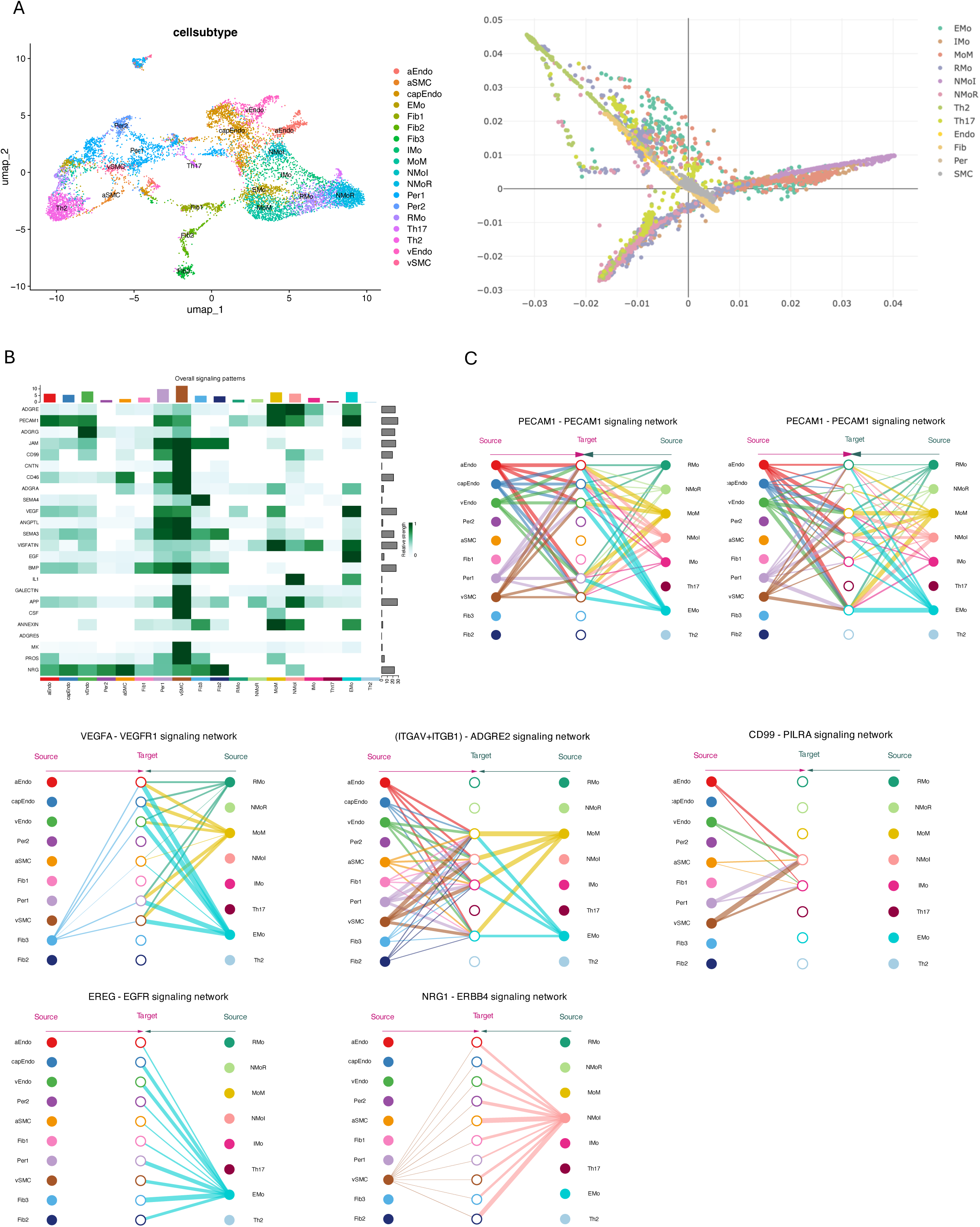
Integration of thrombi snRNA-seq data with publicly available snRNA-seq data from healthy human brain vasculature. (A) Uniform manifold approximation and projection analysis of integrated thrombi and brain vasculature snRNA-seq data. Cells are colored based on cell type. Thrombi cell populations are named according to convention in the rest of the paper (EMo, IMo, MoM, NMoI, NMoR, Th17, Th2). Brain vasculature cell populations are named using metadata from original publication (GEO: 173731). (B) Diffusion component analysis representing diffusion components 1 and 2 for thrombi and brain populations. Here, just as on the UMAP plot, thrombi monocyte populations NMoI and IMo cluster near brain capillary and arterial endothelial cells, thrombi endothelial monocytes cluster near brain endothelial cells and fibroblasts, and Th17 cells cluster near pericytes. (C) Overview of inferred ligand-receptor interactions that may occur between thrombi cell populations and the brain vasculature. (D) Examples of specific ligand-receptor interaction that may occur between specific thrombi cell populations and specific types of brain vasculature cells, which are known to be involved in breakdown of the blood brain barrier, endothelial adhesion, and angiogenesis. Panel 1: PECAM1-PECAM1; Panel 2: VEGFA-VEGFR1; Panel 3: ADGRE-(ITGAV+ITGB1); Panel 4: CD99-PILRA; Panel 5: EREG-EGFR; Panel 6: NRG-ERBB4. aEndo, brain arterial endothelial cells; aSMC, brain arterial smooth muscle cells; capEndo, brain capillary endothelial cells; EMo, thrombi endothelial monocytes; Fib1, brain vasculature fibroblasts cluster 1; Fib2, brain vasculature fibroblasts cluster 2; Fib3, brain vasculature fibroblasts cluster 3; IMo, thrombi inflammatory monocytes; MoM, thrombi monocyte-derived macrophages; NMoI, thrombi neuroepithelial-like inflammatory monocytes, NMoR, thrombi neuroepithelial-like regulatory monocytes; Per1, brain pericytes cluster 1; Per2, brain pericytes cluster 2; RMo, thrombi regulatory monocytes; Th17, thrombi Th17 cells; Th2, thrombi Th2 cells; vEndo, brain venous endothelial cells; vSMC, brain venous smooth muscle cells.

After integration, we used CellChat to infer potential cell type specific ligand-receptor interactions between thrombi cells and the brain vasculature that may occur during stroke (**Fig 3C,D**). Globally, brain vasculature cells engaged in a higher degree of signaling with each other than with thrombi immune cells. However, the volume of potential interactions between certain thrombi cell populations and the brain vasculature approached lower levels of brain-brain signaling. Of all thrombi cell populations, MoM cells engaged in the highest volume of signaling, followed by Th2 cells, then NMoI and IMo cells, which displayed similar levels.

Notable, statistically significant (p<0.05) potential direct cell-cell contact interactions related to leukocyte-endothelial adhesion (*ADGRE-CD55*, *PECAM1-PECAM1*, *ADGRG1-TGM2*, *JAM2-(ITGAV-ITGB1), ADGRE2-(ITGAV-ITGB1), ICAM1-ITGAL*)[69]; transendothelial migration and endothelial activation of leukocytes (both monocytes and T cells) (*NOTCH2-JAG2*, *CD99-PILRA*, *VSIR-IGSF11*, *CNTN1-NOTCH2*, *CD46-JAG1*)[70], [71], [72], [73]; and monocyte extravasation/diapedesis, loss of endothelial and blood brain barrier integrity, and tissue invasion (*SEMA4D-PLXNB2*, *LPAR1-ADGRE5*, *ADGRE5-(ITGAV-ITGB1)*)[74], [75], [76] (**Supp table 2**).

Notable, statistically significant (p<0.05) potential secreted signaling interactions related to angiogenesis, disruption of the blood brain barrier/vessel destabilization, and endothelial and pericyte dysfunction (*VEGFA-VEGFR1*, *VEGFA-KDR*, *VEGFA-(FLT-KDR)*, *ANGPTL2/4-ITGA5_ITGB1*, *SEMA3C-PLXND1*, *NAMPT-(ITGA5-ITGB1)*)[77], [78], [79], [80], [81], [82]; pro-inflammatory pathways (*ACVR1_BMPR2*, *IL1B-IL1R1_IL1RAP*, *APP-TNFRSF21*, *CSF1-CSF1R*, *ANGPTL1-LILRB*, *IL34-CSF1R*)[83], [84]; and anti-inflammatory pathways involving regulation of T cells, complement, the innate immune system, and neuro-immune crosstalk (*ANXA1-FPR3*, *LGALS9_CD45*, *CD55-ADGRE5*, *MDK-LRP1*, *LGALS9_CD44*, *TGFB-TGFBR*)[85], [86], [87], [88] (**Supp table 2**). We also identified two potential interactions related to neural synapse formation, guidance, and plasticity (*NRG1-ERBB4*, *EREG-ERBB4*) [89] (**Supp table 2**).

## Conclusion

We present the first single nuclei level resolution study of the cellular composition of three intracranial human thrombi from stroke patients with large vessel occlusions treated with endovascular thrombectomy. First and foremost, this study demonstrates the feasibility of molecular study of thrombi via single nuclei RNA-sequencing, which is well suited for studying thrombi given the heterogeneous and fragile nature of their tissue. Using snRNA-seq, we identified unique, diverse monocyte and T cell populations in distinct effector states within the thrombus microenvironment, contextualized them in carotid atherosclerotic disease, and inferred potential ligand-receptor interactions that may occur between thrombus cells and the brain vasculature that contribute to inflammation and compromise the blood brain barrier. Using differential gene expression analysis, gene ontology, regulon-level analysis, and integration with carotid plaque data, we provide evidence that monocyte populations in thrombi undergo transdifferentiation and interconversion at the site of occlusion. Finally, we show that inflammatory cell populations in thrombi may interact directly with the brain vasculature and contribute to post-stroke inflammation and microvascular dysfunction at the site of occlusion.

By characterizing the cellular and transcriptional heterogeneity present across thrombi monocyte populations and providing evidence that they undergo transdifferentiation to distinct effector states following embolization due to signals from the local microenvironment, we show that monocytes retain their plasticity in stroke. Monocyte plasticity is well studied in other homeostatic and disease contexts, from tissue repair and angiogenesis, to inflammation and infection; across all of these processes, monocytes are known to undergo rapid transcriptional change to meet the needs of the tissues to which they are recruited [90]. The conservation of monocyte plasticity in stroke not only reflects the dynamic interplay between thrombi and their environment, but also raises the possibility of exploiting this plasticity for therapeutic design. The preservation of monocyte plasticity in stroke also underscores the utility of thrombi as indirect biopsies of the brain, which, with larger scale study, may facilitate the identification of predictive biomarkers and development of novel therapies.

Still, significant challenges remain in elucidating the pathophysiology of post-stroke inflammation and microvascular dysfunction, and connecting these molecular mechanisms to poor outcomes in stroke patients, which our study, designed as a proof of concept for transcriptomic study of thrombi, is not suited in isolation to answer. Going forward, larger scale study of thrombi will be needed to evaluate and characterize the full range of cellular heterogeneity in thrombi, and to mitigate the limitations inherent to all single nuclei or single cell-level study, including processing power, transcript coverage bias, and low capture efficiency, which our study was vulnerable to. Future study must also include wet lab and functional assays to evaluate the molecular significance and/or therapeutic potential of the pathways and cell populations that we identified in this paper, particularly those related to inflammation, loss of endothelial integrity, disruption of the blood brain barrier, and microvascular dysfunction. Finally, incorporating details about patients’ times from their last known normal should also be considered in future analyses involving snRNA-seq to improve understanding of the timeline of post-stroke inflammation on molecular and cellular levels.

Overall, we hope that this study sets the stage for future, larger scale snRNA-seq based study of thrombi, with the end goal of advancing the identification of novel biomarkers for stroke and the development of targeted therapies to reduce inflammation at the site of occlusion in patients who have suffered from acute ischemic stroke.

## Declarations of interest

******

No other author has any decelerations of interest.

## Acknowledgements

The snRNA-seq experiments were conducted at the Single Cell Analysis Core of the Columbia Genome Center and Biomedical Informatics Resources at Columbia University, supported by the NIA T35AG044303 grant.

## Author contributions

E.F.S., V.P.L., P.D.C., M.G.A., P.A.S., and S.E.C. conceptualized the analysis. E.F.S. and V.P.L. performed the analysis, prepared figures, and wrote the manuscript. E.F.S., V.P.L, S.E.C., S.D.L., and G.F.M., retrieved thrombi via endovascular thrombectomy. E.F.S., V.P.L., J.E.A., J.L.F., and C.P.K. performed banking of thrombi specimens. E.F.S., V.P.L., W.Z., A.M., and P.A.S. performed snRNA-seq. All authors edited and reviewed the manuscript.

## Data availability

All single nuclei datasets in this study have been deposited in a Mendeley Data Repository, available via DOI: 10.17632/wt2typ7pzb.1.

## Competing interests

The authors declare no competing interests.

---

**Supplementary Figure 1. Gene set enrichment analyses (GSEAs) reveal unique pathways and phenotypic properties of each thrombi cellular population.**

(A) The Go Biological Process 2023 database reveals upregulation of pathways associated with angiogenesis, vascular smooth muscle regulation, and hypoxia in the EMo cluster.

(B) The WikiPathways 2024 Human database reveals upregulation of pathways associated with neuroinflammation and complement in the IMo cluster.

(C) The WikiPathways 2024 database reveals upregulation of pathways associated with TGFB, T cell receptor signaling, and integrin mediated cell adhesion in the MoM cluster.

(D) The Reactome 2022 database reveals upregulation of pathways related to the adaptive and innate immune systems and complement cascade in the NMoI cluster.

(E) The PanglaoDB 2021 database reveals that NMoR monocytes display a predominately myeloid-like phenotype, rather than a microglial phenotype.

(F) The Go Biological Process 2023 database reveals pathways associated with regulation of phagocytosis and extracellular matrix constituent secretion in the RMo cluster.

(G) The Go Biological Process 2023 database reveals pathways associated with cellular adhesion in the Th2 cluster.

(H) The Go Biological Process 2023 database reveals pathways associated with T cell mediated signaling and T cell differentiation in the Th17 cluster.

EMo, endothelial monocytes; IMo, inflammatory monocytes; MoM, monocyte derived macrophages; RMo, regulatory monocytes; NMoI, neuroepithelial-like inflammatory monocytes; NMoR, neuroepithelial-like regulatory monocytes; Th2, Th2 T cells; Th17, Th17 T cells.

**Supplementary Figure 2. Violin plots depicting expression levels of specific genes that illuminate the unique phenotypic properties of thrombi cellular populations.**

(A) Expression levels of microglia and brain associated genes in the NMoI cluster.

(B) Expression levels of microglia and brain associated genes in the NMoR cluster.

(C) Expression levels of pro-angiogenesis and endothelial adhesion genes in the EMo cluster. EMo, endothelial monocytes; IMo, inflammatory monocytes; MoM, monocyte derived macrophages; RMo, regulatory monocytes; NMoI, neuroepithelial-like inflammatory monocytes; NMoR, neuroepithelial-like regulatory monocytes; Th2, Th2 T cells; Th17, Th17 T cells.

**Supplementary Figure 3. Seurat’s Azimuth reference based pipeline cell type identifications**

UMAP plot depicting thrombi cells with identity predictions from Seurat’s Azimuth mapping algorithm supports our cell type identifications.

**Supplementary Figure 4. SCENIC analysis identifies differential regulon-level activity across all thrombi cellular populations**

SCENIC regulon analysis reveals differential regulon activity among all thrombi cell populations. Each row represents a specific transcription factor and each column represents an individual cluster. The dendrogram at the top represents relatedness among thrombi cell populations on the basis of regulon activity. Activity is relative, with red representing higher levels of activity for a given transcription factor, and blue representing relatively lower levels of activity.

EMo, endothelial monocytes; IMo, inflammatory monocytes; MoM, monocyte derived macrophages; RMo, regulatory monocytes; NMoI, neuroepithelial-like inflammatory monocytes; NMoR, neuroepithelial-like regulatory monocytes; Th2, Th2 T cells; Th17, Th17 T cells.

**Supplementary Figure 5. Thrombi cells express complement related genes**

Violin plot demonstrating expression of complement receptors and complement factors across thrombi cell populations.

EMo, endothelial monocytes; IMo, inflammatory monocytes; MoM, monocyte derived macrophages; RMo, regulatory monocytes; NMoI, neuroepithelial-like inflammatory monocytes; NMoR, neuroepithelial-like regulatory monocytes; Th2, Th2 T cells; Th17, Th17 T cells.

## Notes

### Competing Interest Statement

The authors have declared no competing interest.

