## Supplementary figures and images for "THE MICROANATOMY OF INTRACRANIAL HUMAN THROMBI BY SINGLE NUCLEI TRANSCRIPTOMICS"

### Supplemental Figure 1

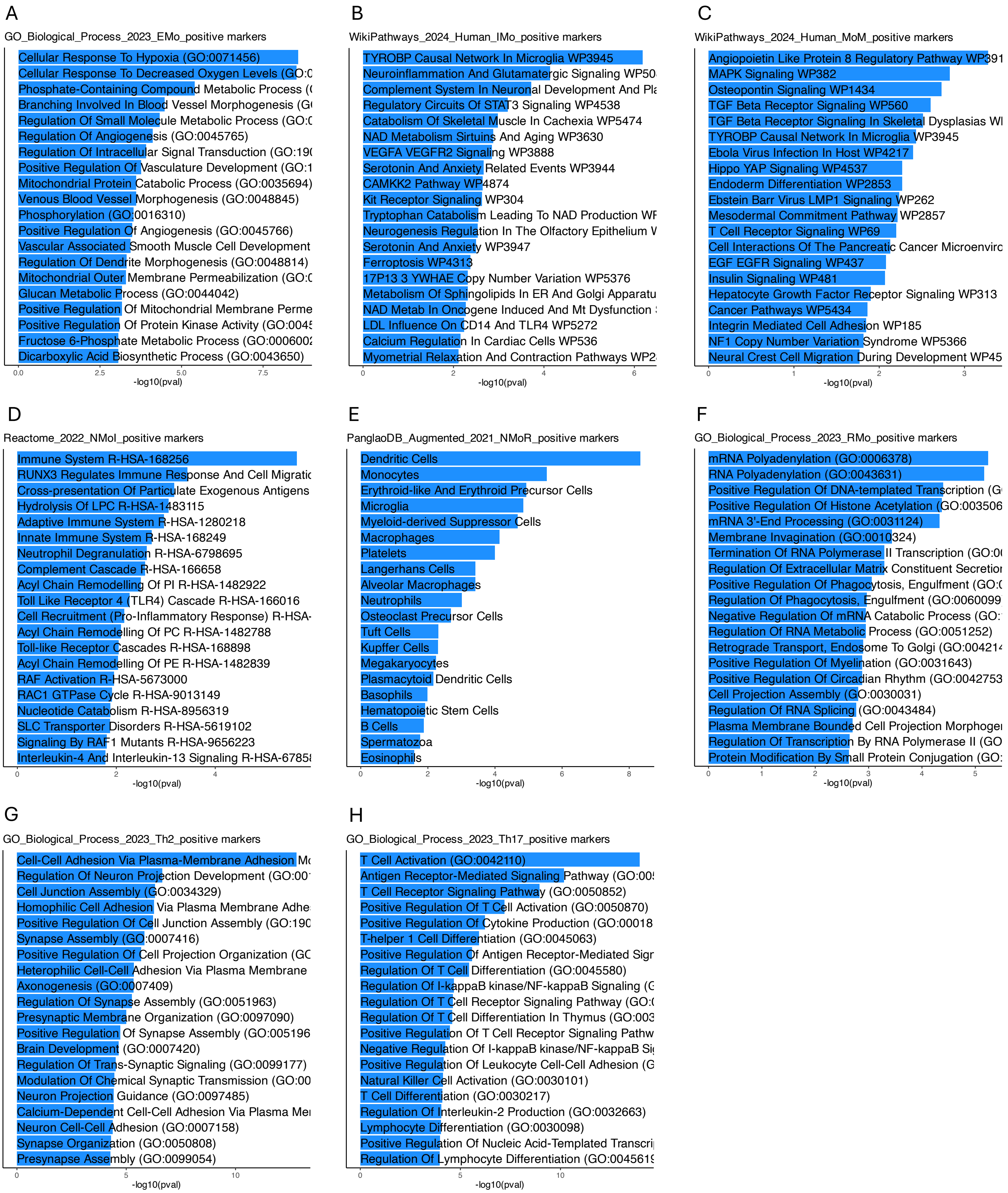


Supplemental Figure 1

### Supplemental Figure 2

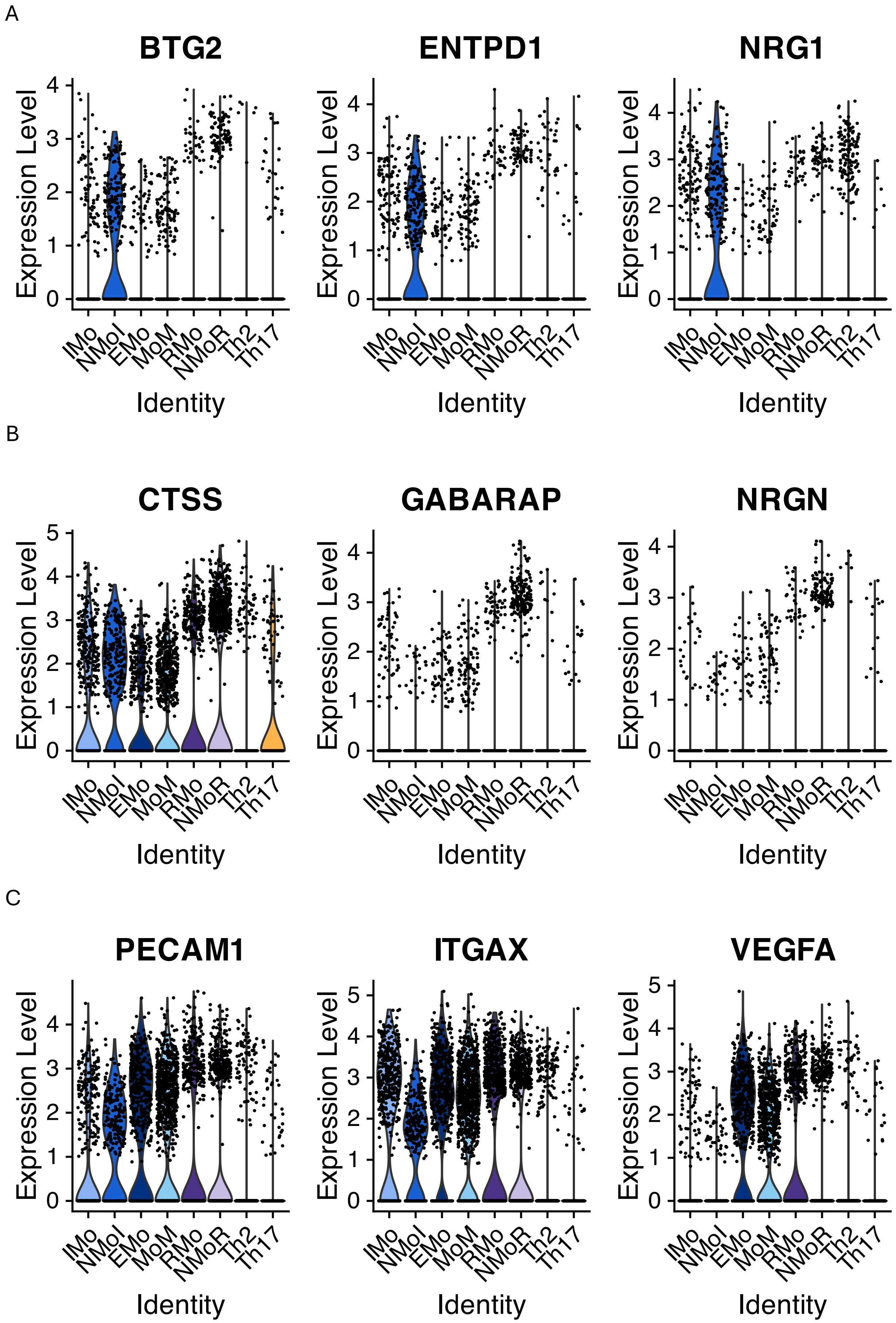


Supplemental Figure 2

### Supplemental Figure 3-4

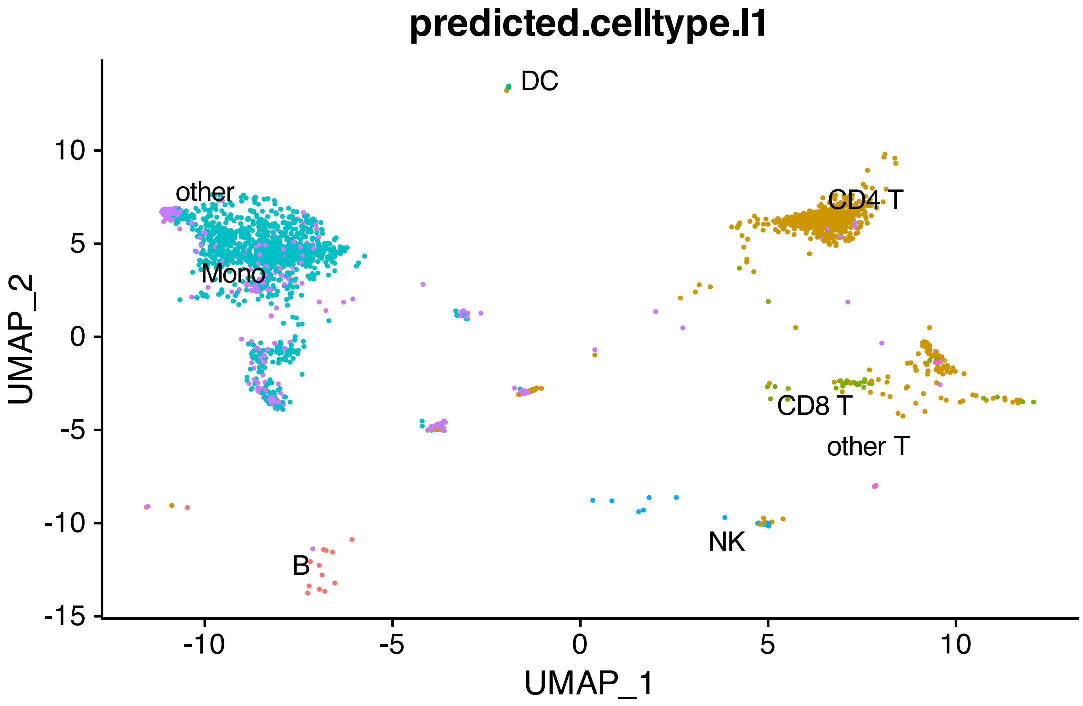

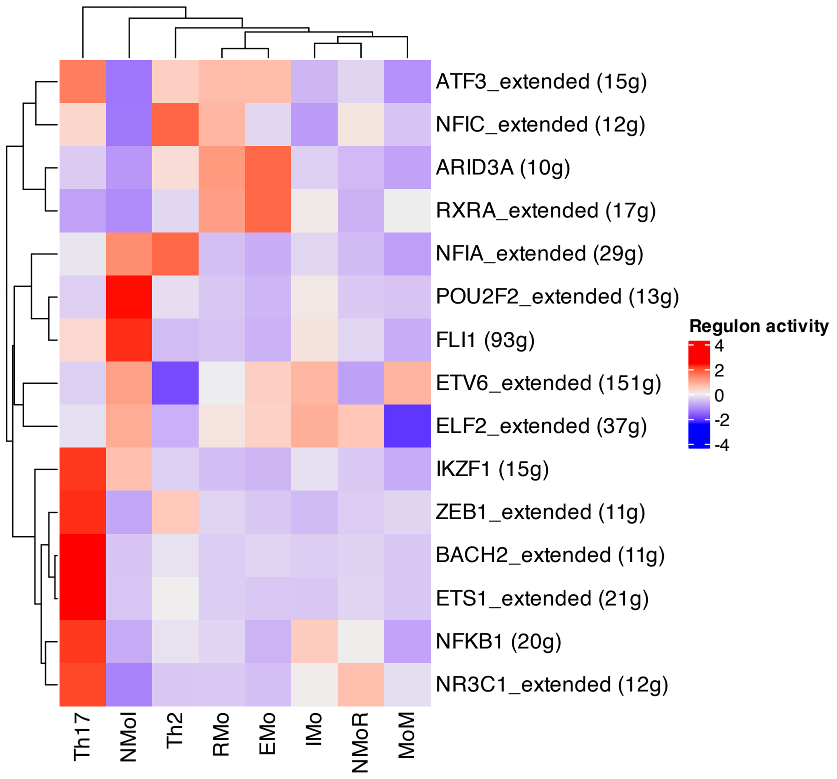

### Supplemental Figure 5

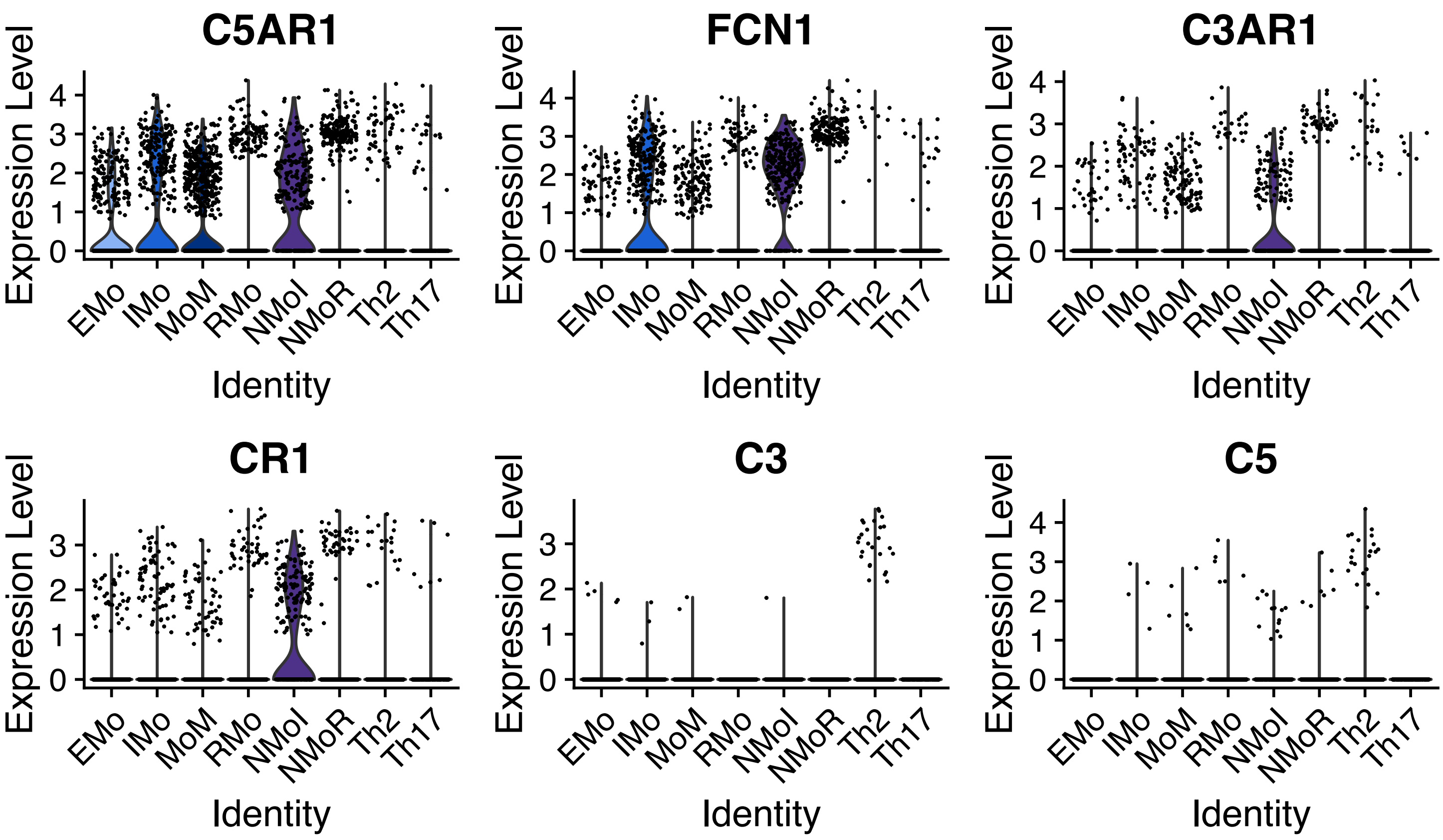


Supplemental Figure 5
